# Listening with your heart: The heartbeat shapes auditory object formation by suppressing the early neural response to sound

**DOI:** 10.64898/2026.08.17.745255

**Authors:** John P. Veillette, Aditi Joshi, Ying Li, Howard C. Nusbaum

## Abstract

Interoceptive sensations, arising from the body’s visceral organs such as the heart, are known to impact exteroceptive sensory perception. The classical explanation for heart-to-brain influence, the Baroreceptor Hypothesis, posits that baroreceptors firing during the systolic blood pressure peaks that follow each heartbeat suppress the magnitude of neural responses to exteroceptive sensations. More recent work, however, has demonstrated qualitative (rather than merely magnitude) differences in perception as a function of the cardiac cycle; since it is not obvious how the Baroreceptor Hypothesis could explain these findings, even in principle, they have often been characterized as incompatible. We propose baroreceptor-related suppression of early sensorineural responses need not manifest simply as suppression of corresponding conscious percepts. In a validated computational model of auditory cortex that segregates an ambiguous tone sequence into either one or two auditory objects or “streams,” we found suppression of the neural response to one tone type increases the likelihood that tone is parsed into a distinct stream. We subsequently verified this prediction empirically: when presenting such ambiguous sequences to human participants in a manner such that one tone type only occurs during cardiac systole, the initial neural response to that tone—indexed by the electroencephalographic (EEG) frequency-following response (FFR)—is indeed suppressed, while participants report hearing the sequence as two separate sounds streams more frequently. Thus, suppression of early sensorineural responses can be sufficient to explain qualitative, not just magnitude, differences in perception when considered in the context of larger neural circuits.

**Significance Statement:** Interactions between the heart and brain have been proposed to underlie our subjective experience of ourselves and of the world, conditioning conscious perception on each individuals’ internal state. For these interactions to meaningfully affect our experience, however, would require a biological mechanism by which sensory inputs from the body alter the categorical content of experience, not just superficial aspects of perception (e.g., perceived magnitude). Seemingly in contrast, the dominant theory of heart-brain interaction posits that cardiac input merely reduces neural responses to external sensations. We show, through computational modeling and empirical experimentation, that simply suppressing the magnitude of the neural response to sound can indeed alter the content of perception, influencing how we parse an acoustic scene into perceptual objects.

## Introduction

Interoceptive sensory inputs arising from the body’s visceral organs, such as the heart, routinely impact how the brain processes sensory signals from the external environment (Critchley and Harrison, 2013; Engelen et al., 2023). Lacey and Lacey were among the first to note that responses to exteroceptive stimuli differ by time of presentation within the cardiac cycle, presumably because aortic baroreceptors—the primary sensory transducers in the heart-to-brain axis—fire during ventricular systole, when blood pressure peaks, while remaining relatively silent during ventricular diastole (Lacey and Lacey, 1970, 1978). Their Baroreceptor Hypothesis, that baroceptor input to the brain has a broadly inhibitory effect on exteroceptive sensory function, remains highly influential (Azzalini et al., 2019).

Consistent with the Baroreceptor Hypothesis, exteroceptive stimuli presented coincident with baroreceptor activity are less likely to be detected and are perceived as less intense across modalities including vision (Salomon et al., 2016, 2018), audition (Schulz et al., 2009), and nociception (Dworkin et al., 1994; Wilkinson et al., 2013). Beyond simple psychophysical measurements, things get more complicated. For example, visual discrimination (Pramme et al., 2014) and search (Pramme et al., 2016) can be facilitated during ventricular systole relative to diastole. Perception of some emotional stimuli also appears facilitated (Gray et al., 2012; Garfinkel et al., 2014). The Baroreceptor Hypothesis seems to make no clear prediction at all about other findings. During systole, for example, faces are perceived as more trustworthy (Azevedo et al., 2023), sociocultural stereotypes are more potent (Azevedo et al., 2017), and time intervals are perceived as shorter (Arslanova et al., 2023). Stimuli synchronized to systole are more likely to be experienced as part of oneself, as in the rubber hand illusion (Aspell et al., 2013; Suzuki et al., 2013; Sel et al., 2017). Such findings demonstrate the heart can alter the high-level contents of conscious awareness, not just whether or how strongly something is perceived. This motivates the idea that interoceptive-exteroceptive interactions, by conditioning the content of conscious awareness on our bodies’ internal states, are responsible for salient aspects of subjective experience: from the first person perspective of conscious awareness (Park and Tallon-Baudry, 2014; Azzalini et al., 2019) to the sense of a unified, embodied self (Seth and Tsakiris, 2018).

It is not obvious the Baroreceptor Hypothesis, positing only inhibition of sensory cortical function by baroreceptors, could possibly explain findings at higher levels of psychological organization than simple psychophysical relationships (e.g., perceived magnitude). We argue it might still. Though Lacey and Lacey seemed to favor a more psychological definition of “inhibitory” where cardioceptive inputs pull some finite processing resource like attention away from exteroception (Lacey and Lacey, 1974), their hypothesis has since come to be understood as implying actual neural inhibition to suppress temporally predictable sensory consequences of the heartbeat (Seth, 2013; van Elk et al., 2014; Salomon et al., 2016, 2018), analogous to inhibitory corollary discharge from motor to sensory cortices during voluntary movements (Von Holst and Mittelstaedt, 1950; Blakemore et al., 2000). False equivocation between these two interpretations makes baroreceptor inhibition seem incompatible with many of the above findings, but in the context of larger neural circuits, inhibition of sensorineural activity need not result in suppression of the corresponding conscious percept. For example, Wilkinson et al. (2013) speculate that baroreceptor-suppression of cortical activity releases the spinal cord from cortical inhibition, explaining facilitated somatosensory detection during systole (Wilkinson et al., 2013).

In the present study, we first incorporated Baroreceptor-Hypothesis-like inhibition into an existing neural population model of an auditory stream segregation task, where participants are presented with an ambiguous tone sequence that is bistably perceived as coming from either one or two source streams—a discrete and qualitative, rather than just a magnitude, difference in perception. We used this model to show that manipulating the timing of tone presentation around baroreceptor activity should impact stream perception, and we subsequently confirmed this behavioral prediction—and the prediction of sensorineural suppression—in human participants.

## Materials and Methods

### Participant recruitment and ethics

We recruited University of Chicago students to participate in our study through the Psychology Department’s research participation system. Any student without any previously diagnosed or suspected hearing loss was eligible to participate. We aimed to recruit 20 participants, which was determined to provide adequate statistical power (see *Sample size determination*). However, we could not achieve skin-to-electrode impedances for electroencephalography (EEG) recordings below 10 KΩ in two participants, more than the recommended maximum for our signal of interest (Zhao et al., 2019; Kerneis et al., 2023), so these participants were excluded and replaced—resulting in 22 total participants recruited, though only *n*=20 were analyzed. Of those *n*=20, 12 were biologically male and 8 female. 18 were right-handed, and 2 were left-handed. The average age of participants was 19.4 years, and ages ranged between 18 and 21 years. All procedures were approved by the Social and Behavioral Sciences Institutional Review Board at the University of Chicago (protocol IRB24-1904).

### Bistable auditory stream segregation paradigm and its computational model

Auditory scene analysis is the process by which the brain decomposes the overlapping mixture of sound waves transduced in the ear into meaningful auditory objects or “streams” corresponding to individual sound sources (Bregman, 1994). The auditory stream segregation paradigm, in which participants classify sequences of two or more repeating tones as coming from one or more streams, has been the dominant experimental model for auditory scene analysis for decades (Bregman and Campbell, 1971). In the present study, we consider what is likely the most common variation of this paradigm, introduced in van Noorden’s (1975) influential doctoral dissertation. Here, high (A) and low (B) tones are presented in a repeating ABA-pattern, where the hyphen indicates a rest, and tone durations are one half of the interstimulus asynchrony (see Fig. 1a). When the frequency difference (Δ*f*) between A tones and B tones is small, participants experience ABA-sequences as a single, galloping rhythm (like ABA-ABA- and so on). When Δ*f* is sufficiently high, participants experience the sequence as two metronome-like rhythms playing simultaneously (like A-A-A-A- and -B---B--). Critically, across a relatively broad range of intermediate Δ*f*s, participants may experience the tone sequence as coming from either one or two streams in a mutually exclusive manner (see Fig. 1a), rather than perceiving it as an intermediate mixture between those two possible percepts (van Noorden, 1975). In other words, conscious perception of the ABA-sequence is bistable, as in visual binocular rivalry (Carmel et al., 2010), discretely perceiving one or the other percept at any given time. Usefully for our present purposes, however, likelihood of a participant perceiving a given discrete percept is influenced by certain continuous stimulus parameters, such as Δ*f* or tone amplitude—making this a powerful paradigm for investigating how attenuation of the low-level neural response to sensation affects downstream determinants of perceptual content. Whether a participant perceives the “integrated percept” (one stream) or the “streaming percept” (two streams) is usually assessed via self-report, as we do in the present experiment, but more objective behavioral assays have been used to verify that integrated and streaming percepts are indeed qualitatively distinct and mutually exclusive; for example, participants can only report on the relative timing of A and B tones when they experience the integrated percept (Snyder and Alain, 2007).

**Figure 1:**
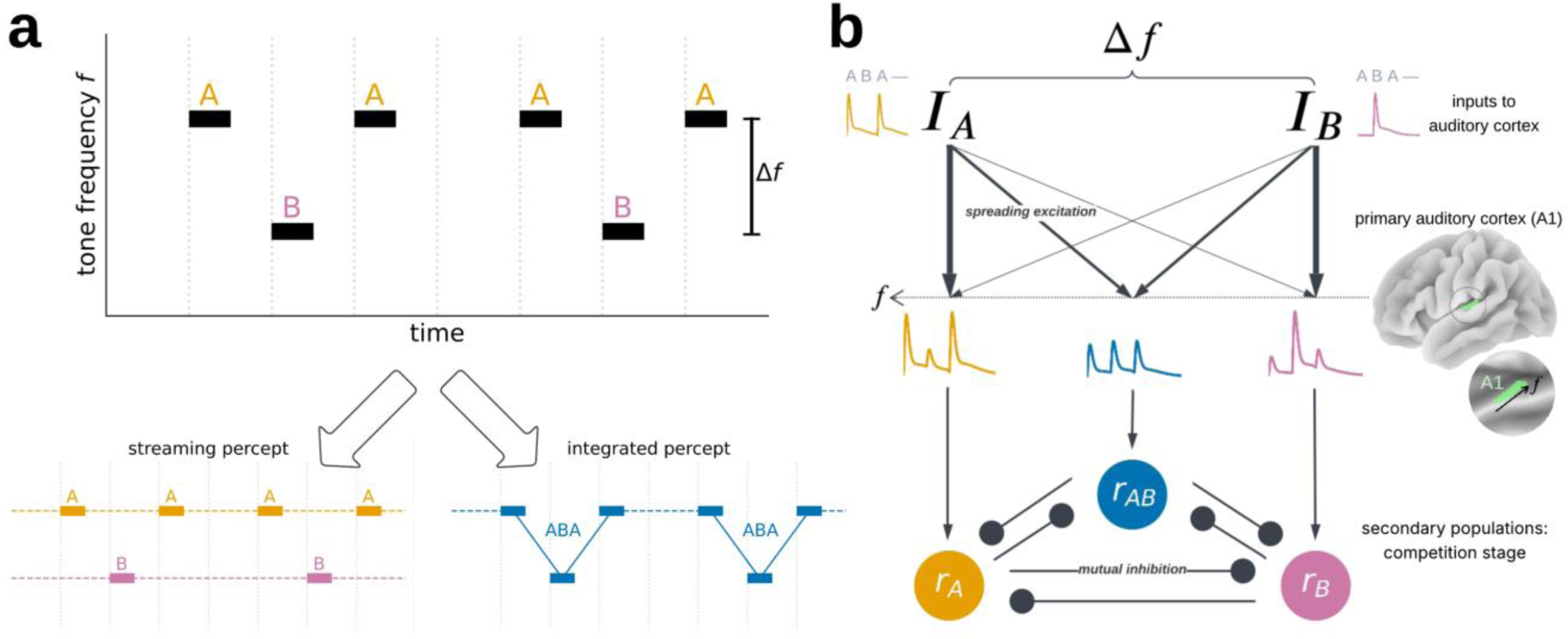
The auditory stream segregation paradigm and contemporary theory. (a) In a common lab paradigm for auditory scene analysis, the process by which we decompose the complex sound wave entering our ear into independent sound sources or “streams,” ABA tone triplets are played repeatedly, separated by a rest. For a wide range of frequency differences Δ*f* between the A and B tones, they are experienced as either one of two mutually exclusive percepts: a “streaming” percept in which participants hear two, metronome-like rhythms playing simultaneously, or an “integrated” percept where participants hear a single, galloping rhythm. (b) This bistable perceptual phenomenon is thought to occur because spatially spreading activation along the tonotopic gradient of primary auditory cortex, where spatially nearby neurons respond maximally to sounds of similar frequencies, means that there is often overlap in the set of neurons responding to both tone types. Contemporary models of auditory stream segregation posit that downstream populations, excited by those primary auditory cortex responses, engage in a mutually inhibitory competition. When a population that responds similarly strongly to both A and B tones dominates that competition, successfully suppressing A- and B-selective populations, one hears the integrated percept. When then A- and/or B-selective populations dominate, one instead hears the streaming percept.

A common account of auditory streaming supposes that the phenomenon is explained by the basic organization of the auditory system, which up to and including primary auditory cortex (A1) is tonotopically organized. That is to say that spatially adjacent neurons in A1 maximally respond to similar acoustic frequencies (Saenz and Langers, 2014). However, neurons exhibit a “tuning curve” by which they still respond albeit with less intensity to sounds nearby their characteristic frequency (Recanzone et al., 2000; Bitterman et al., 2008). Thus, auditory cortex exhibits spreading activation, such that it can be assumed that between A1 areas selective for the frequencies of A tones and of B tones, neurons may respond to both tones (see Fig. 1). When the frequency difference Δ*f* is large enough, however, auditory cortex responses to A and to B tones will be sufficiently separated in tonotopic space that few cells respond to both tones, segregating their downstream neural representations from one another (Fishman et al., 2001; Bee and Klump, 2004). Notably, however, the degree of tonotopic overlap may be determined not just by Δ*f* itself but also by the strength of spreading activation (see Fig. 2c)—exactly what the Baroreceptor Hypothesis claims should be modulated by cardiac phase.

**Figure 2:**
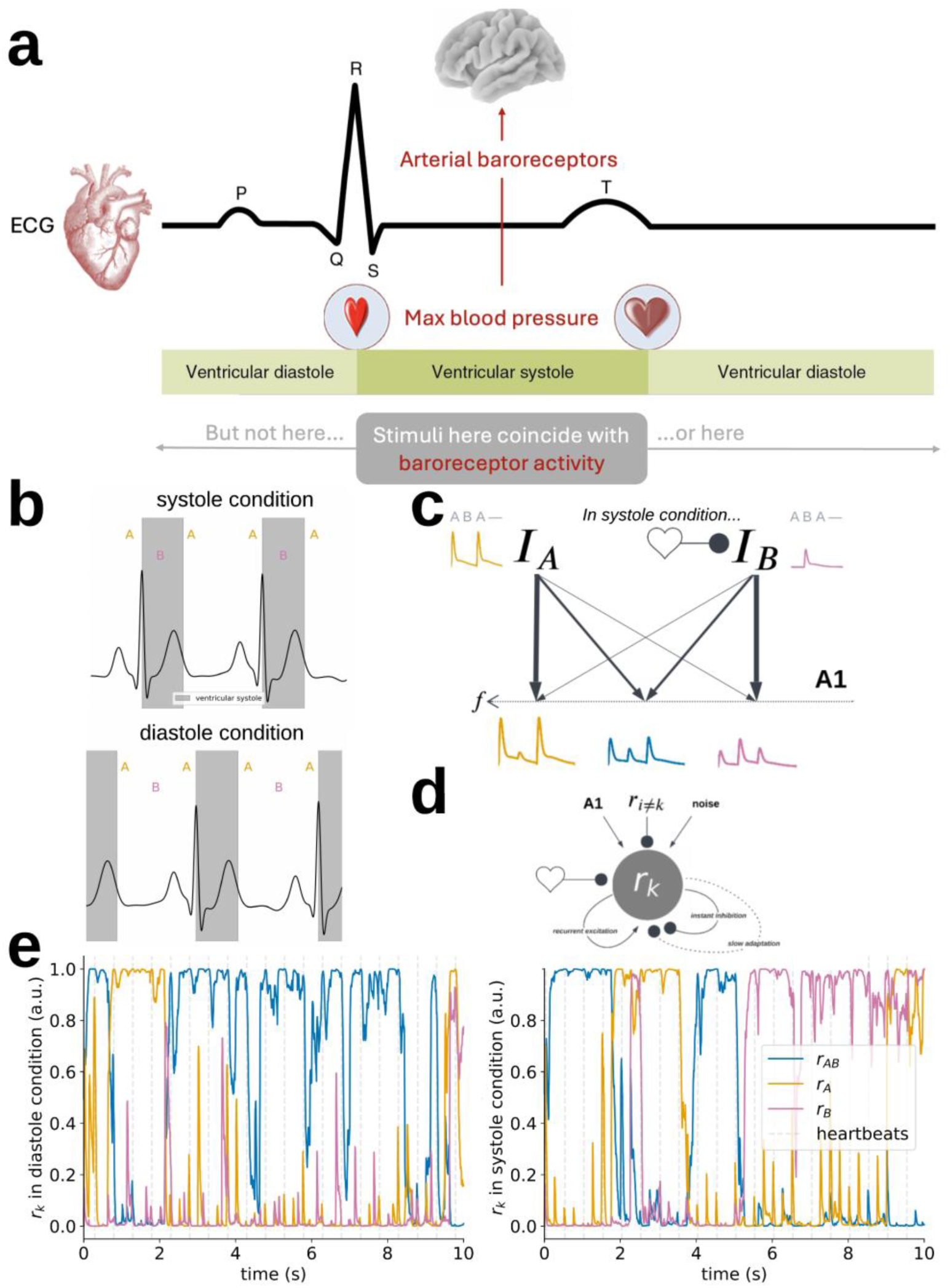
Simulating the effect of cardiac afferent input on auditory stream segregation. (a) Each time the heart contracts, the electrical initiation of which is marked by the QRS complex in the electrocardiogram (ECG) waveform, it ejects newly oxygenated blood into the aorta, causing a transient increase in blood pressure that persists until the heart’s ventricles relax at the end of ventricular systole. This systolic blood pressure peak causes aortic baroreceptors, essentially pressure sensors, to fire—signaling to the brain that a heartbeat has occurred. (b) We consider an auditory streaming paradigm in which the B tones are timed to onset at approximately in the middle of ventricular systole in the “systole condition” and during diastole in the “diastole condition.” Thid cardiac phase manipulation ensures that the B tone coincides baroreceptor input to the brain only in the systole condition. (c) The influential Baroreceptor Hypothesis predicts that baroreceptor input should suppress the neural response to the B tone. However, this suppression would not just affect the maximally B-selective area of auditory but also those areas that respond to both A and B tones. This state of affairs makes it difficult to intuit how baroreceptor-related suppression would alter stream perception, if at all. (d) To make a directional prediction, we simulate the effect of the cardiac phase manipulation on a well-established computational model of those mutually inhibitory cortical populations from Figure 1b during auditory stream segregation (Rankin et al., 2015). (e) Simulations of the average firing rates of mutually inhibitory populations during one task trial, with identical initial conditions and noise processes but different cardiac phase conditions, are shown for illustration. Across thousands of such simulations, we found that presenting B tones coincident with baroreceptor input increased the probability model would “perceive” the streaming percept (see Fig. 1) on a given trial by roughly 18%.

Based on observations from auditory cortical recordings in humans and nonhuman primates during auditory stream segregation paradigms, Rankin et al.’s (2015) neuromechanistic model of bistable stream perception formalizes this conceptual description. The model assumes ascending sensory inputs corresponding to the A tones (*I_A_*) and B tones (*I_B_*) are tonotopically mapped onto primary auditory cortex (A1) as illustrated in Figure 1b (Rankin et al., 2015). *I_A_*(*t*) and *I_B_*(*t*) are the standard Heaviside function that mimics recorded responses in nonhuman primate recordings, convolved with the tone onsets. An A-selective area responds primarily to the A tone, specifically according to *D_A_*(*t*) = *I_A_*(*t*) + *w*(Δ*f*)*I_B_*(*t*) where *w*(·) is a function describing the decay of neural responses with increasing tonotopic distance from the maximally responsive location (Rankin et al., 2015). AB-selective area responds according to *D_B_*(*t*) = *I_B_*(*t*) + *w*(Δ*f*)*I_A_*(*t*), and a hypothetical middle population responds equally to both tones according to 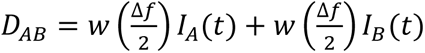 (see Fig. 1b). These primary A1 responses excite downstream populations *r_A_*, *r_B_*, and *r_AB_*, which engage in a mutually inhibitory competition that determines the outcome of perception. They are modeled as a dynamical system, with temporal evolution governed by a system of stochastic differential equations that capture known properties of neural populations:

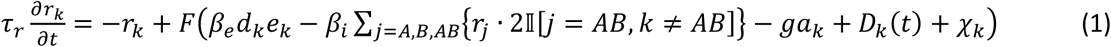

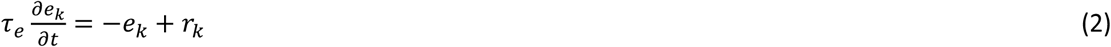

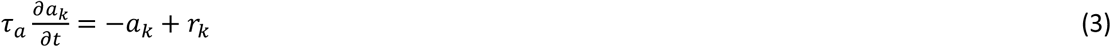

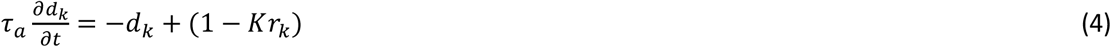

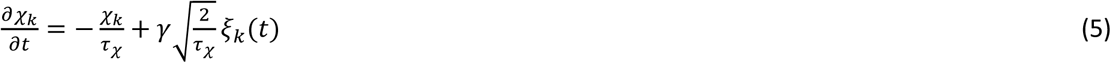

where Equation 2 reflects recurrent excitation, the −*β_i_* ∑*_j_*_=*A*,*B*,*AB*_{*r_j_* · 2‖[*j* = *AB*, *k* ≠ *AB*]} term in Equation 1 reflects self and lateral inhibition which is stronger going from *r_AB_* to *r_A_* and *r_B_* than in the other direction by a factor of two (when indicator variable ‖[*j* = *AB*, *k* ≠ *AB*] is equal to 1). Equation 3 reflects spike frequency adaptation, Equation 4 reflects synaptic depression, and Equation 5 is an Ornstien-Uhlenbeck noise process integrating over white noise *ξ_k_*(*t*). Constant *τ* parameters reflect the characteristic timescales of these different mechanisms. Numerical values of constant parameters, function definitions of sigmoidal activation function *F*(·) and exogenous inputs *I*(·), and detailed justifications of the model specification are originally given in Rankin et al. (2015); however, please note that values of some of the model’s parameters are misreported in that original paper, which the authors corrected in a subsequent manuscript (Byrne et al., 2019).

If one assumes that the simulated person experiences the integrated percept when *r_AB_* > 0.5(*r_A_* + *r_B_*) and the streaming percept otherwise, this model reproduces the empirically observed statistics of spontaneous switching between integrated and streaming percepts over seconds an minutes in healthy humans (Rankin et al., 2015) as well as cochlear implant patients (Paredes-Gallardo et al., 2019). It has also been used successfully to predict, *a priori*, the effects of novel stimulus manipulations on stream perception (Byrne et al., 2019), as we aimed to do in the work described here.

### Incorporating cardioceptive input into model of auditory stream segregation

We considered a “systole condition,” where the onset of the B tones always followed the heartbeat thus falling during ventricular systole and coincided with baroreceptor activity (see Fig. 2a, and a “diastole condition” where systole occurred after the last A tone of each triplet such that all tones were presented during diastole (see Fig. 2b). We simulated 10 second ABA-sequences with an interstimulus asynchrony of 1/8 of a second (i.e. one whole triplet occurring every half second) and Δ*f* = 5 semitones, the frequency difference at which Rankin et al. (2015) reported approximate equidominance (i.e., maximal ambiguity) between percepts. The neural response to the heartbeat *H*(*t*) was simulated, like to inputs to auditory cortex evoked by the A and B tones (Rankin et al., 2015), as a Heaviside function convolved with the onset of the hypothetical heartbeats but with four times the width as the responses to the tones, such that modeled systole could encompass a full tone presentation. In line with the modern Baroreceptor Hypothesis, the neural response to the heartbeat was treated as a source of global inhibition to auditory cortex (see Fig. 2d). It was incorporated into a modified version of Equation 1, as follows in Equation 6:

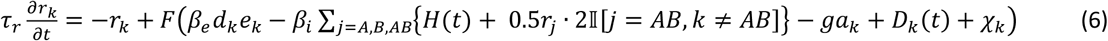

In the systole condition, the onset of the heartbeat was simulated as occurring 1/12 of a second before the onset of the B tone. In the diastole condition, it was delayed by 1/4 of a second relative to in the systole condition, such that the simulated neural response to the heartbeat occurred during the rest between ABA triplets (see Fig. 2b). Additionally, the N1 component of the auditory-evoked EEG response—classically thought to include (though not specific to) the low-order response of primary auditory cortex to sound (Woods, 1995)—is known to be reduced by cardiac input (van Elk et al., 2014). Since the modifications in Equation 6 only affect the downstream *r_k_* populations, presumably located in a secondary auditory area (Rankin et al., 2015), we also divided the amplitude of *I_B_* by half only in the systole condition to reflect reduced ascending drive to auditory cortex (see Fig. 2c). The choice to divide by half, and not by some other value, was arbitrary—but the goal of this exercise was simply to generate a directional prediction that we could test in an empirical experiment, rather than to maximize biological realism.

We simulated *r_A_*, *r_B_*, and *r_AB_* firing rates for 1000, 10-second trials of each condition by numerically integrating differential Equations 2-6 over time using the Julia programming language’s *DifferentialEquations.jl* package (Rackauckas and Nie, 2017a, 2017b). Simulations were paired across conditions, such that the first simulation of the systole condition used the same random seed and initial conditions as the first simulation of the diastole condition, and so on; one such paired trial is visualized in Figure 2e. We assumed, following the original model (Rankin et al., 2015), the simulated participant would “report” hearing the integrated percept if *r_AB_* > 0.5(*r_A_* + *r_B_*) for the majority of the trial, and they would report hearing the streaming percept if 0.5(*r_A_* + *r_B_*) > *r_AB_* for the majority of the trial. Under these assumptions, the simulated participant labeled 46.2% of the diastole trials as integrated but only 39.1% of systole trials, indicating that baroreceptor-related suppression during the neural response to the B tone should have the effect of facilitating the streaming percept. All model simulations were performed prior to collecting any empirical data. Subsequently, we performed an experiment with human participants to test the model’s directional prediction.

### Experimental design: Auditory stream segregation task

Each participant listened to 100, 10-second trials of repeating ABA-tone sequences, as in our model simulations. We recorded ECG and detected occurrences of the ECG R-peaks (see Fig. 2a) in real time (see *Electrocardiography (ECG) recording and online R-peak detection*), around which we timed the presentation of the tones. To recreate the conditions from our simulations, the interstimulus onset asynchrony and tone durations had to be set based on the online ECG on a trial-by-trial basis, given between- and within-participant variation in heart rates. At the start of each trial, we calculated the median inter-beat interval (IBI, i.e., duration between consecutive R-peaks) of the 10 preceding heart beats. Tone durations for that trial would be set to 1/8 of the IBI and interstimulus intervals to 1/4 of the IBI. In the systole condition, B tones were triggered by each subsequently detected R-peak to onset (1 + 1/6)*IBI seconds later, or approximately 1/6 of the IBI after the following R-peak—roughly the middle of ventricular systole. The A tones were triggered by each subsequently detected R-peak to onset (1/6 + 3/4)*IBI and (1/6 + 5/4)*IBI later, respectively preceding the following R-peak and trailing the subsequent systole. On diastole trials, the B tones were instead triggered to onset during diastole at (1/6 + 3/2)*IBI following each detected R-peak—directly antiphase to their position on systole trials—while A tones continued to fall at the same cardiac phase (see Fig. 2b). After each trial, participants were asked whether they heard a single, galloping rhythm (coherence) or two, metronome rhythms (streaming). Since perception is bistable and participant were expected to switch between the two percepts spontaneously over the course of the trial (Snyder and Alain, 2007), just as our model tends to (see Fig. 2e), we specifically asked participants which percept they had experienced more by the end of the trial. Participants were not told about the cardiac phase manipulation in advance; instead, they were told that we were recording ECG because we were interested in whether their perception influences their heart rate (i.e., the opposite causal direction from that in which we were actually interested).

The experiment took place in a soundproof Faraday cage. All tones were played to participants over Etymotic Research 3C insert earphones, which physically isolate the participant from the electrical sound generator by directing pressure to the earphone through a rubber tube to avoid contaminating EEG recordings. Tones were generated using the *PsychToolbox* driver via *PsychoPy* (Brainard and Vision, 1997; Peirce, 2007; Peirce et al., 2019), were pure sine waves windowed with a 5 ms Hanning taper at onset and offset to reduce clipping artifacts during payback, and they were played at an amplitude of 65-70 dB SPL, verified with a sound level meter. B tones were always of frequency 500 Hz. Since the range of frequency differences Δ*f* at which perception of ABA-sequences is bistable depends on the interstimulus interval (Snyder and Alain, 2007), which was determined in real time, it was not possible to set a single Δ*f* that was constant across all participants and trials. Instead, the frequency of the A tones was assigned on a trial-by-trial basis by a Bayesian adaptive QUEST+ routine, which fit a psychometric function (predicting integrated/streaming responses as a function of stimulus frequency) after each trial and chose the next frequency in the range 510-5000 Hz based on a minimum entropy selection rule—in other words, it picked as the next A tone frequency that at which it was most uncertain about how the participant would response, ensuring Δ*f* was always in the range where perception of the ABA-was maximally ambiguous (Watson, 2017). 50 trials were randomly assigned to the systole condition and 50 to the diastole condition; randomization was performed on a participant-by-participant basis such that no participants received the conditions in the same order. This randomization affords causal inference for the effect of systole/diastole condition assignment, manipulating the coincidence of baroreceptor activity with the B tone (see Fig. 2a-b), on stream segregation judgments for the ambiguous ABA-sequences.

### Statistical analysis: Auditory stream segregation task

The effect of systole/diastole condition assignment on integrated/streaming judgments was quantified using a mixed-effects logistic regression with a fixed effect for condition a random intercept for each participant. Condition was coded as 1 for systole and 0 for diastole, and participant judgments were coded as 1 for streaming and 0 for integrated. Regressions were fit using R’s *lme4* v2.0.1 package through its *pymer4* Python interface (Bates et al., 2015; Jolly, 2018). We report the fixed effect coefficient and its parametric 95% confidence interval on its native log-odds scale, and we report non-parametric bootstrap 95% confidence intervals for the marginal mean proportion of streaming responses in each condition and for the condition difference, computed using *lme4*’s *bootMer* function. All bootstrap confidence intervals reported in text are “highest density intervals,” the smallest interval that contains 95% of bootstrap samples.

We opt to report only mean/parameter estimates and 95% confidence intervals, in accordance with recent recommendations to emphasize the statistical uncertainty surrounding measured effects rather than merely the binary inference implied by *p*-values (Calin-Jageman and Cumming, 2019a, 2019b; Ho et al., 2019). However, we note the equivalence between a 95% confidence interval excluding zero and the a two-tailed *p*-value being less than 0.05; thus, results presented here do still use the typical statistical significance threshold (i.e., false positive rate) of *α* = 0.05 for inference.

### Sample size determination

We estimated our statistical power to detect the predicted behavioral effect with a bootstrap power calculation, using the output of our model simulation (Liu, 2026). In other words, for each bootstrap simulation, we resampled 50 x *n* systole and 50 x *n* diastole trials—that is, 100 trials for each simulated participant—with replacement from the 1,000 trials we had simulated of each condition (see *Incorporating cardioceptive input into model of auditory stream segregation*). We then fit a mixed-effects logistic regression, as in our planned analysis (see *Statistical analysis: Auditory stream segregation task*), to the resampled data. We repeated this procedure 1,000 times, recording whether the confidence interval of the fixed effect coefficient for condition contained zero. Power was estimated as the proportion of simulations in which the 95% confidence interval excluded zero; thus, this estimate is of statistical power at a false positive rate of *α* = 0.05. This calculation determined that statistical power of 90% (or, more specifically, 89.8%) was achieved with *n*=20, which we used as our sample size.

### Experimental design: Heartbeat discrimination task

One alternative explanation for any effect of manipulating stimulus timing within the cardiac cycle is always that, if participants are able to perceive their own heartbeat, they become aware of the cardiac timing manipulation and direct their attention to the stimulus differently as a result. To assess this possibility, all participants completed a modified version of the heartbeat discrimination task after they completed the main streaming task to measure their interoceptive awareness (Whitehead et al., 1977). In a heartbeat discrimination task, participants are asked to judge whether a stimulus is synchronous to or delayed relative to the sensation of their heartbeat. Because there are substantial individual differences in when people perceive their heartbeat sensations (Brener and Kluvitse, 1988; Brener et al., 1993), it is critical when using heartbeat discrimination as a control task to match the cardiac phase manipulation closely across the two tasks—such that performance on the heartbeat discrimination task corresponds to participants’ ability to discriminate between the experimental conditions in the main task (Veillette et al., 2024).

Thus, on each trial, 500 Hz “B tones” were played with the same two possible cardiac phase onsets and same durations as they would be in the main task (see *Experimental design: Auditory stream segregation task*), this time in the absence of A tones. There were again 100 trials, with 50 randomly assigned to have a systole onset and 50 randomly assigned to a diastole onset. This time, after each 10-second trial, participants were explicitly asked to judge whether they thought the tones played in synchrony with or delayed relative to their heartbeats.

### Statistical Analysis: Heartbeat discrimination task

In explicit interoceptive awareness tasks like heartbeat discrimination (Brener and Kluvitse, 1988; Brener et al., 1993) and in implicit coupling between cardioception and other modalities (Engelen et al., 2025), it is a common finding that while a majority of participants may show a cardiac timing effect at some cardiac phase of stimulus presentation, there are substantial differences in which phase that is—such that less than 30% of participants may show an effect for a given, fixed cardiac phase contrast. While this fact may be inconvenient from the perspective of designing well-powered experimental studies, it can also be quite convenient for inference in the sense that every cardiac timing manipulation has a built-in quasi-experimental control. To assess whether a cardiac timing effect on exteroceptive perception requires participants to be consciously aware of their own heartbeat, one can simply check whether some participants who are unable to explicitly report on the cardiac phase manipulation in the heartbeat discrimination control task still show an effect of that manipulation in the main task (Veillette et al., 2024). In other words, the scientific question “Does this effect require awareness of the heartbeat?” becomes a population prevalence question: “Are there people in the population who show this effect but are not able to explicitly report on their heartbeat?” This sort of prevalence inference can be accomplished using Bayesian mixture modeling (Ince et al., 2021, 2022; Veillette and Nusbaum, 2025).

To estimate the joint population prevalence of the auditory streaming effect and heartbeat discrimination sensitivity—that is, the joint probability that a participant randomly drawn from the population would show one effect, the other, or both—we use a within-group *p*-curve mixture model as specified and validated in our previous methods paper (Veillette and Nusbaum, 2025). In brief, we calculate a *p*-value (used not for null hypothesis significance testing but rather as a convenient transformation of variables) for each task and individual participant, using a logistic regression of response vs. condition as in our main analysis (see *Statistical analysis: Auditory stream segregation task*). A *p*-curve mixture model treats the population distribution of *p*-values as a mixture between a uniform distribution for participants who do not show an effect and a right-skewed curve, parametrized with a single *Z*-scale size parameter per effect of interest, for participants who do show the effect. A within-group *p*-curve mixture model, in particular, estimates mixture parameters corresponding to the set of probabilities ℙ[*H*_1_ is true, *H*_2_ is false], ℙ[*H*_1_ is true, *H*_2_ is true], ℙ[*H*_1_ is false, *H*_2_ is false], and ℙ[*H*_1_ is false, *H*_2_ is true] for a random person drawn from the sampled population. From these quantities, one can also calculate marginal prevalence estimates like ℙ[*H*_1_ is true] as well as conditional estimates like ℙ[*H*_1_ is true|*H*_2_ is false].

We report posterior means and 95% highest-density credible intervals for the prevalences of an auditory stream segregation effect and of above-chance interoceptive awareness. We then report the conditional prevalence of the stream segregation effect in the subpopulation who is not aware of the cardiac phase manipulation. Posterior distributions for model parameters were approximated with 5,000 posterior samples generated using Markov Chain Monte Carlo (Hoffman and Gelman, 2014), and we verified that the convergence metric *r*^ = 1.0 for all parameters, indicating good model convergence.

To specifically quantify evidence that a stream segregation effect either does or does not require awareness of the cardiac timing manipulation, we report a Bayes Factor comparing the full within-group *p*-curve mixture model to a nested model without the ℙ[*H*_1_ is true, *H*_2_ is false] parameter, where *H*_1_ is the streaming effect and *H*_2_ is awareness of the cardiac timing manipulation. Bayes Factors are computed using bridge sampling (Gronau et al., 2017). All *p*-curve mixture models are implemented with our open source *p2prev* Python package, which is a wrapper around the *PyMC* probabilistic programming language (Patil et al., 2010; Veillette and Nusbaum, 2025).

For completeness, we also report the bootstrap 95% confidence interval for the group mean interoceptive awareness, i.e. heartbeat discrimination sensitivity quantified as *d*-prime (Sumner and Sumner, 2020).

### Electroencephalography (EEG) recording and preprocessing

In order to test the direct prediction of the Baroreceptor Hypothesis—that the initial neural response to the B tone, corresponding to the input to primary auditory cortex *I_B_* in our model, is suppressed when stimulus presentation coincides with baroreceptor activity in ventricular systole—we also recorded EEG throughout the experiment. We placed a recording electrode at Fz (approximate hairline), the reference electrode on the left mastoid, and the ground electrode on the right mastoid to measure the frequency-following response (FFR), a non-invasive measure of neural firing phase-locked to the acoustic stimulus waveform in the ascending auditory system (Moushegian et al., 1973), which has sources beginning at the auditory nerve and extending all the way into (tonotopic) primary auditory cortex (Behroozmand et al., 2016; Coffey et al., 2019). The electrode placement was selected to optimally record the neural response to sound ascending from auditory midbrain structures into the auditory cortex, based on its parallel alignment to vertical dipole of the auditory brainstem (Bidelman, 2015), though we do not claim that this dipole is the only contributing source (Coffey et al., 2019). In other words, the FFR is as direct a measure as can be obtained non-invasively of the ascending drive *I_B_* to tonotopic primary auditory cortex; moreover, as a frequency-tagged measure, it has the practical benefit of being far outside of the range of ECG artifacts in the EEG trace or of overlapping heartbeat-evoked potentials, the presence of which might otherwise be confounded with the cardiac timing condition (Park and Blanke, 2019). We used Ag–AgCl ring electrodes, and voltages were recorded using an ActiCHamp Plus amplifier (Brain Products, GmbH) and an EP-PreAmp (Brain Products, GmbH) bipolar preamplifier with a gain of 50. EEG signals were digitized at a sampling rate of 20,000 Hz, and the ground truth onsets of tone presentations were marked in the recording using a StimTrak (Brain Products, GmbH) hardware sound level trigger through which participants’ earphones were passed with an Acoustical Stimulator Adapter.

The magnitude of the frequency-following response to a pure tone stimulus is indexed by spectral power—specifically at the frequency of the acoustic stimulus—in the evoked potential obtained by averaging over hundreds or thousands of repetitions of that tone (Krizman and Kraus, 2019). To obtain this quantity, we first notch filtered the data at the line noise frequency (60 Hz) and its harmonics up to 2,000 Hz, and then we applied a bandpass filter with a pass band from 100-2,000 Hz. For each participant, we took their minimum tone duration *D* (since tone durations were set on a trial-by-trial basis, see *Experimental design: Auditory stream segregation task*) and collected epochs of -D to +D seconds around the onsets of all B tones presented across both tasks. We did not analyze EEG data from the tasks separately due to the large number of stimulus repetitions recommended to obtain clean FFR waveforms (Krizman and Kraus, 2019). We subtracted the mean voltage from the baseline of -D to 0 seconds, removed any trial where to peak-to-peak amplitude exceeded a predefined rejection threshold of 35 μV. This resulted in, on average, 989 epochs in the systole condition and 961 epochs in the diastole condition remaining after artifact rejection. We then averaged over all remaining epochs within each condition to obtain a time-domain evoked waveform for each condition. We computed the power spectral density of the Hanning-tapered pre-stimulus baseline and post-stimulus evoked EEG for each condition using multitaper analysis (Slepian and Pollak, 1961), and we finally quantified evoked spectral power in the FFR relative to baseline power in decibels. All EEG preprocessing was done using the *MNE-Python* package (Gramfort et al., 2014).

### Statistical analysis: EEG frequency-following response (FFR)

We report non-parametric bootstrap 95% confidence intervals for evoked FFR power at the 500 Hz frequency of the B tone when it is presented during systole and during diastole, and we also report the 95% confidence interval for the paired difference between those conditions.

### Electrocardiography (ECG) recording and online R-peak detection

We recorded ECG onboard the same EEG amplifier using a Lead II electrode configuration, and we accessed ECG data for online processing using the Lab Streaming Layer framework (Kothe et al., 2025). ECG signals were to decimated from the recorded 20,000 Hz to 100 Hz and then bandpass filtered to 5-15 Hz before R-peaks were detected using the online Pan-Tompkins algorithm (Pan and Tompkins, 1985). Developed for our previous work (Veillette et al., 2024), our implementation of the Pan-Tompkins algorithm is a port of Sznajder and Łukowska’s (2017) to Meta Reality Labs’s open *LabGraph* software (https://github.com/facebookresearch/labgraph) for easy interoperability with *PsychoPy* and other Python-based software.

### Data and code availability

All raw behavioral and EEG/ECG data, as well as preprocessed FFRs, are available on *OpenNeuro* (Markiewicz et al., 2021) at https://doi.org/10.18112/openneuro.ds007949.v1.0.0. Data are organized and documented according to the standardized Brain Imaging Data Structure to ensure long-term interpretability and usability (Gorgolewski et al., 2016). Model simulation code, which remains unaltered since prior to the beginning of data collection, is permanently archived on *Zenodo* at https://doi.org/10.5281/zenodo.21432897; this archive also contains code for our sample size calculation. The code used to run the empirical experiments (including ECG R-peak detection) is archived at https://doi.org/10.5281/zenodo.21432881, and electrophysiology preprocessing and all analysis code is archived at https://doi.org/10.5281/zenodo.21432866.

## Results

### The initial neural encoding of sound is suppressed by the heartbeat

When B tones were presented during ventricular systole, when blood pressure is high and aortic baroreceptors fire (see Fig. 2a), the response of the ascending auditory system to that tone was suppressed relative to when it was presented during diastole (see Fig. 3a). During diastole, the evoked power of the EEG frequency-following response was 13.94 dB (95% CI: [11.61, 16.11]). During systole, it was only 12.95 dB (95% CI: [10.72, 15.12]), or −0.9886 dB less (95% CI: [−1.91, - 0.11]).

**Figure 3:**
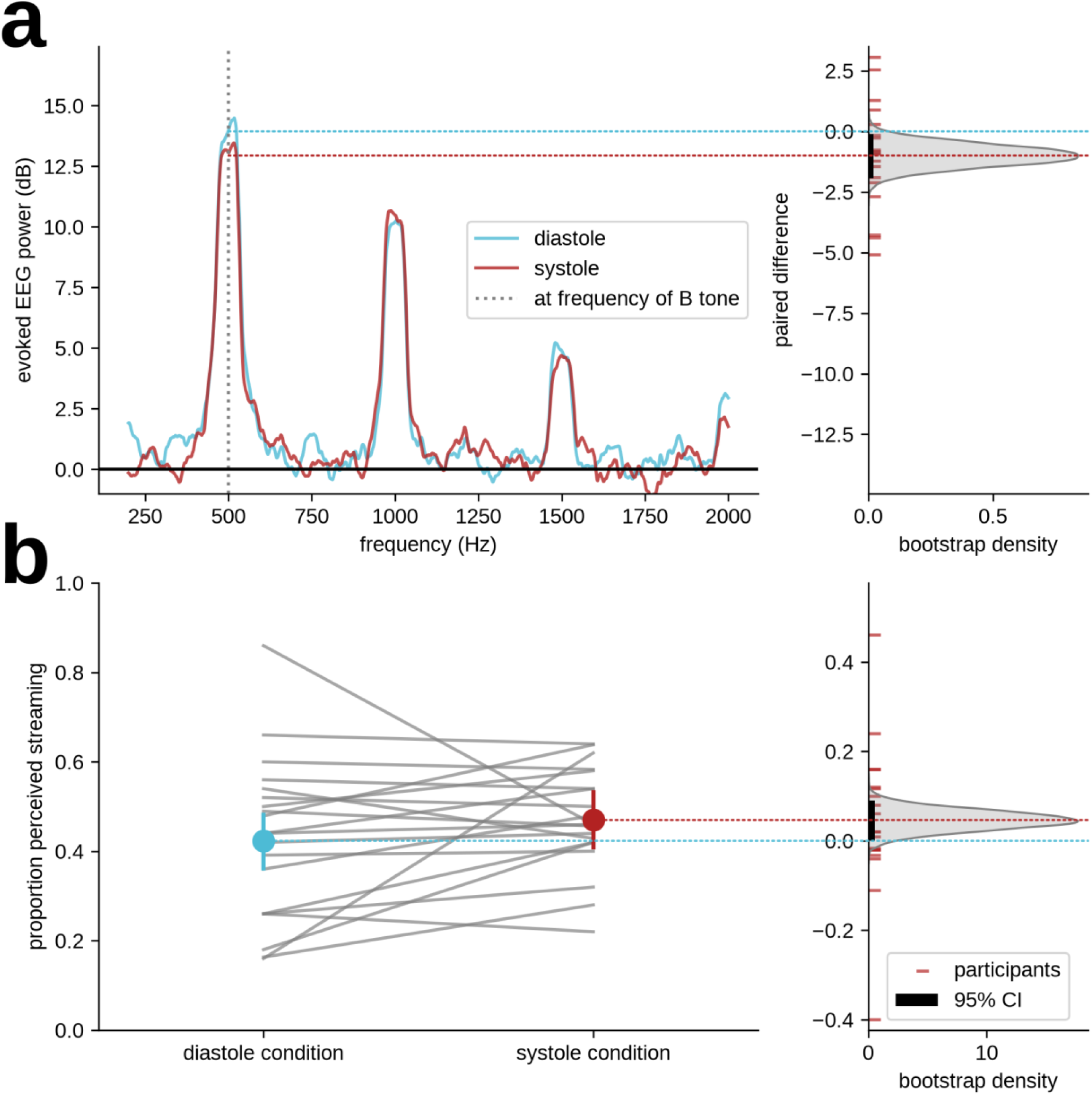
Empirical validation that cardiac afferent input suppresses the neural response to tones while facilitating perceived segregation of sounds into multiple streams. (a) The EEG frequency-following response is a non-invasive measure of neural firing phase-locked to the periodic oscillation of an acoustic stimulus, a hallmark of the ascending auditory system up to and including primary auditory cortex. The amplitude of this stimulus-evoked response in human participants is suppressed when the stimulus is presented during systole relative to when it is presented during diastole. Evoked EEG power spectral densities relative to the pre-stimulus baseline is shown on the left, while the bootstrap distribution of the paired mean difference between conditions is shown on the right. (b) As the computational model predicted would be the effect of such sensorineural suppression, the streaming percept—that is, hearing the ABA-sequence as two, simultaneous metronome like rhythms rather than as one, integrated rhythm— is more likely to be experienced when B tones are presented during systole. Individual participants’ rates of streaming are shown on the left, and the bootstrap distribution of the marginal mean difference is again shown on the right.

### Baroreceptor input synchronous to the B tone facilitates its perception as a separate stream

When ambiguous ABA- tone sequences were presented such that the B tone was always coincident with the expected time of baroreceptor activity, participants were more likely to experience the ABA-ABA-sequences as two separate metronome-like streams (like A-A-A-A- and -B---B--, see Fig. 1a) than they were when all tones in the triplet were presented during ventricular diastole, when baroreceptors should be relatively silent (see Fig. 2a). On average, participants reported that they heard the two-stream percept on 42.33% (95% CI: [36.14, 48.19]) of trials in the diastole condition but 46.99% (95% CI: [40.85, 53.21]) during the systole condition. Per a mixed-effect logistic regression, this is a β=0.189 (95% CI: [0.007, 0.372]) log-odds increase in likelihood of experiencing the streaming percept from diastole to systole, or an absolute difference in percentage of 4.67 (95% CI: [0.15, 9.10]) as visualized in Figure 3b.

### Baroreceptor effect on auditory stream segregation does not require awareness of the heartbeat

On average, participants were able to report on when tones were presented relative to their cardiac cycle with slightly above-chance sensitivity (*d’*=0.27, 95% CI: [0.12, 0.50]). Consistent with past literature, however, population prevalence inference showed that only a modest fraction of people in the population are, in fact, able to discriminate between cardiac phase conditions at all (posterior mean=0.29, 95% CI [0.08, 0.54]). The effect of baroreceptor input on stream segregation, as expected while presenting stimuli at a fixed cardiac phase in each condition (see *Statistical Analysis: Heartbeat discrimination task*), was also estimated to be present in only a fraction of the population (posterior mean=0.22, 95% CI: [0.05, 0.42]). Importantly, the expected prevalence of the streaming effect among people who cannot explicitly discriminate between the cardiac timing conditions is essentially unchanged from that among the full population (posterior mean=0.22, 95% CI: [0.01, 0.45]). In fact, comparing the full population model to a nested model in which participants cannot show the streaming effect unless they can explicitly discriminate between the cardiac phase conditions yielded a Bayes Factor of BF=8.75 in favor of the full model. In other words, the observed data are 8.75 times more likely to occur in a world where the baroreceptor effect on auditory stream segregation does *not* require participants to be consciously aware of the timing of the stimuli relative to their heartbeats than they are to occur in a world where it does.

## Discussion

Consistent with the prediction of the Baroreceptor Hypothesis, the present work demonstrates that the ascending neural response of the early auditory system is suppressed when a sound is heard during ventricular systole, the window of peak baroreceptor activation, relative to when it is heard during diastole. Simultaneously, an auditory stimulus that might have been grouped with its auditory background was more likely perceived as a distinct auditory object when accompanied by synchronous baroreceptor activation. This latter effect constitutes a qualitative/categorical difference in auditory scene perception. However, by simulating the effect of our experimental manipulation in an existing neural population model of stream segregation in auditory cortex, we show that inhibition of sensorineural responses alone is sufficient to account for this qualitive difference—no additional mechanism required. We see this finding as a potential first step toward explaining the complex and varied effects of cardiac afferent input on high-level subjective experiences from biological mechanism up. For example, given that the exteroceptive sensations most likely to synchronize with the heart in practice are those that result from events in the body itself (Balakrishnan et al., 2013; Gaardbaek and Kidmose, 2025), the fact that such sensations are more likely to be parsed into distinct perceptual objects may facilitate known effects of cardiac synchrony on perceived embodiment (Aspell et al., 2013; Suzuki et al., 2013; Sel et al., 2017).

This sufficiency of sensorineural suppression to produce a qualitative difference in perception, rather than simply a difference in perceived magnitude as one might intuitively predict, highlights a logical flaw in how sensory suppression is often discussed in human neuroscience. As reviewed in the Introduction, cardiac afferent input has been observed to affect superficial social judgments from faces (Azevedo et al., 2017, 2023), emotion perception (Gray et al., 2012; Garfinkel et al., 2014), time interval perception (Arslanova et al., 2023), putative implicit measures of the sense of agency (Herman and Tsakiris, 2020; Koreki et al., 2022), as well as self/other discrimination and subjective embodiment (Aspell et al., 2013; Suzuki et al., 2013; Sel et al., 2017). Since such qualitative effects on subjective experience do not intuitively follow from the Baroreceptor Hypothesis, however, there is a strong temptation to explain them instead by other mechanisms.

Evidence for the existence of other plausible mechanisms by which interoceptive rhythms such as the heartbeat can influence brain processes, as reviewed recently (Engelen et al., 2023), is reasonably strong. For example, the magnitude of the evoked response to the heartbeat covaries with visual detection thresholds (Park et al., 2014) and bodily self-consciousness (Park et al., 2016) even when stimuli are not presented in systole. In all likelihood, the inhibitory mechanism predicted by the Baroreceptor Hypothesis coexists with other interacting mechanisms, like interoceptive modulation of effective functional connectivity in the brain (Kim and Jeong, 2019). It would be a mistake, however, to prematurely assign a given perceptual or behavioral interoceptive-exteroceptive interaction effect to one of these other mechanisms only based on a false equivalence between neural and perceptual suppression under the Baroreceptor Hypothesis. The present findings, while demonstrating neural suppression, are arguably inconsistent with perceptual suppression or Lacey and Lacey’s early supposition that baroreceptor input draws attention away from exteroceptive modalities (Lacey and Lacey, 1974), as reducing attention directed toward a given tone type should reduce the likelihood that tone is segregated into a separate auditory stream (Sussman et al., 1999).

Instead, our results are consistent with a modern interpretation of baroreceptor-related sensory suppression as analogous to the inhibitory corollary discharge from motor to sensory cortices during voluntary movement (Seth, 2013; van Elk et al., 2014; Salomon et al., 2016, 2018). Interestingly, this analogy extends to the aforementioned false equivocation between neural and perceptual suppression. Though corollary discharge has long been thought to uniformly reduce perceived stimulus magnitude, a recent series of studies has found exteroceptive perception during movement appears facilitated in some contexts (Yon and Press, 2017; D’Onofrio Pacheco and Zimmermann, 2026), though this claim has been contested (Job and Kilteni, 2023). Such findings have been characterized as challenging longstanding dogma in motor control (Press et al., 2023). Similar to interoceptive influences on embodied self-awareness, attenuation of the neural response to sensorimotor feedback is also often—though not always (Timm et al., 2016)— accompanied by hallmark qualities of subjective experience that do not intuitively follow from suppression, such as the sense of agency (Veillette et al., 2023, 2025).

Rather than construing directionally heterogenous of qualitative perceptual/behavioral consequences of cardiac afferent input—or of internal motor signals for that matter—as incompatible with sensory suppression, we argue it is critical to consider inhibition of early sensorineural responses in the context of larger neural circuits and systems. While some authors have noted baroreceptor-related cortical suppression could plausibly manifest as perceptual facilitation in some contexts (Wilkinson et al., 2013), work that considers how Baroreceptor-Hypothesis-predicted effects on early sensorineural responses should alter the operating characteristics of well-defined neural computations downstream of those initial responses is almost nonexistent. The little work that does exist, however, highlights the utility of computational models for predicting the complex, often unintuitive consequences of parsimonious cardiac modulation of sensory responses (Allen et al., 2022).

As employed in the present work, bistable perceptual phenomena—in which participants experience only one of two or more mutually exclusive percepts—may be uniquely productive experimental models for how continuous changes in the strength of low-level sensorineural responses can produce discrete, qualitive changes in perception (Safavi et al., 2026). Such paradigms produce large subjective differences in perception while still affording strong experimental control, and seemingly minor design variations can reveal substantial nuance in the machinery of interoceptive-exteroceptive integration. In interocular rivalry, for example, the relative strength of the brain’s responses to different visual stimuli presented to each eye is understood to determine which of those binocularly presented stimuli occupies conscious awareness (Carmel et al., 2010). When one “masking” stimulus is substantially higher contrast than a “masked” stimulus presented to the opposite eye—a continuous flash suppression paradigm (CFS)—presenting the masked stimulus during ventricular systole prevents it from breaking into awareness, nominally consistent with the Baroreceptor Hypothesis (Salomon et al., 2016). When instead the binocularly presented stimuli are more evenly matched—a typical binocular rivalry paradigm— we have found a transient stimulus manipulation known to increase stimulus strength (Wade et al., 1984) is more effective during systole than during diastole (Veillette et al., 2024). In other words, the directional effect of the same cardiac phase manipulation depends on the extant strength relationship between rival stimuli, as both stimuli are presented simultaneously, and the effect of baroreceptor input is likely non-selective over the ecologically-meaningless stimuli used by both Salomon et al. (2016) and Veillette et al. (2024). (In contrast, stimuli in the presently bistable auditory stream segregation paradigm are presented over time, affording selective baroreceptor modulation of the response to one tone, as illustrated in Figure 2a-b, though rival percepts here do not correspond to the different tone types but rather how they could be parsed into inferred sound sources.) It was subsequently shown, even in a CFS paradigm, masked stimuli can be facilitated by baroreceptor input when it is emotionally salient (Ozturk et al., 2025). Interoceptive influences on visual bistability, like the auditory bistability investigated here, are ripe for more concrete computational specification to generate empirically testable predictions that will refine our understanding of underlying biological mechanisms (Yonekura et al., 2025).

The widespread cardiac influence on subjective experiences of ourselves and of the world have generated much excitement that interoceptive senses could hold the secret to understanding conscious self-awareness (Park and Tallon-Baudry, 2014; Seth and Tsakiris, 2018), with potential applications to a wide range of clinical mental health conditions in which interoception is known to be anomalous (Khalsa et al., 2018). Any such account, however, is incomplete without a well-defined biological mechanism by which interoceptive inputs can alter the qualitative content, not just superficial aspects, of perception. The present work highlights the importance of combining empirical experimentation with circuit and system level computational modeling to bridge the gap between specific neural phenomena and their often unintuitive consequences in perception. The half century old Baroreceptor Hypothesis, though even we once declared its demise (Veillette et al., 2024), may prove to have more life in it yet.

## Author Contributions (CRediT Taxonomy)

**Conceptualization:** J.P.V.; **Data curation:** J.P.V.; **Formal analysis:** J.P.V., A.J., and Y.L.; **Funding acquisition:** H.C.N.; **Investigation:** A.J. and Y.L.; **Methodology:** J.P.V.; **Project administration:** J.P.V., A.J., and Y.L.; **Resources:** H.C.N.; **Software:** J.P.V.; **Supervision:** J.P.V. and H.C.N.; **Validation:** J.P.V.; **Visualization:** J.P.V.; **Writing - original draft:** J.P.V.; **Writing - review & editing:** J.P.V., A.J., Y.L., and H.C.N.

## Acknowledgements

This work used research equipment that was paid for by a grant from the National Science Foundation (NSF BCS 2024923) awarded to H.C.N.

## Conflict of Interest Statement

The authors declare no conflicts of interest.

## Notes

https://doi.org/10.18112/openneuro.ds007949.v1.0.0

https://doi.org/10.5281/zenodo.21432897

https://doi.org/10.5281/zenodo.21432881

https://doi.org/10.5281/zenodo.21432866

### Competing Interest Statement

The authors have declared no competing interest.

